# Pulmonary macrophage subsets display distinct metabolic responses to polarising stimuli *in vivo*

**DOI:** 10.1101/2025.07.02.662727

**Authors:** Joshua J. Hughes, Stefano A.P. Colombo, Lee Booty, Amanda J. L. Ridley, Benjamin Harrop, Grace Mallet, Rafael J. Argüello, Alexander Phythian-Adams, Andrew S. MacDonald

**Author notes:** Corresponding author: Professor Andrew. S. MacDonald.

## Abstract

Macrophage activation is underpinned by metabolic changes required to fight infection, resolve inflammation and enable effective wound healing. While metabolic control of macrophage activation is increasingly understood in culture systems *in vitro*, it remains poorly understood in more complex *in vivo* settings, like the lung. Here we applied novel flow cytometry based immunometabolic techniques to profile immune cell metabolism in the murine lung. We revealed a surprising role for glucose in naïve alveolar macrophages (AMs) that was retained by AMs polarised *in vivo* with either interleukin-4 (IL-4) or lipopolysaccharide (LPS). We identified that naïve interstitial macrophages (IMs) were dependent on mitochondrial and glucose metabolism, IMs polarised with IL-4 failed to induce metabolic alterations but displayed a glycolytic phenotype following LPS exposure as they adopted an ‘M1’ like metabolic profile. We also demonstrated that AMs were metabolically less responsive than IMs to intranasal delivery of LPS, but upregulated glycolysis and metabolic features of ‘M2’ polarisation (defined *in vitro*) in response to intranasal IL-4, including oxidative phosphorylation (OXPHOS), fatty acid oxidation (FAO) and arginine metabolism. Finally, we identified AM M2 polarisation as highly sensitive to glucose inhibition *ex vivo*. Thus, lung macrophage subsets display distinct metabolic responses to polarising stimuli *in vivo*.

**Highlights:**

- Naïve alveolar macrophages require glucose metabolism despite residing in a low glucose environment.
- Alveolar macrophages are more responsive to IL-4 *in vivo* than LPS upregulating oxidative metabolism, lipid metabolism and glycolysis.
- Interstitial macrophages adopt a glycolytic phenotype characteristic of ‘M1’ BMDMs *in vitro* following *in vivo* LPS administration.
- Alternatively activated alveolar macrophages are extremely sensitive to glucose inhibition *ex vivo*.

## Introduction

During early embryogenesis, macrophages populate our tissues performing organ-specific functions required to maintain health^1–4^. Tissue-specific macrophage phenotype and function is shaped by the local environment. One mechanism by which this is regulated is the control of available metabolites which, in the lung, is primarily mediated by the epithelium^5–9^. Surfactant, a lipid rich fluid, is secreted into the airway to maintain efficient gas exchange^10–12^. Without proper regulation surfactant accumulates, leading to respiratory failure, and the primary role of resident airway alveolar macrophages (AMs) is to phagocytose and degrade excess surfactant^13, 14^. Although lipids are abundant, glucose is rapidly transported out of the airways, restricting pathogen growth and creating a uniquely low glucose environment^10–12^. AMs are therefore thought to rely on oxidative phosphorylation (OXPHOS) to meet bioenergetic requirements and display a blunted glycolytic metabolism^15^. The tightly regulated metabolic environment of AMs does not extend to lung tissue resident interstitial macrophages (IMs), which reside in glucose concentrations over 12-fold greater than the airway^16^. Although a role for glycolysis in IM activation has been suggested in the context of bacterial infection^17, 18^, their metabolism remains poorly characterised, particularly in the absence of exogenous activation. Overall, the metabolic phenotype of pulmonary macrophage subsets in homeostasis or following activation *in vivo* remains unclear and requires further investigation.

As the dominant immune population in the airway, AMs act as a first line of defence against damage to the airway epithelium and pathogen invasion ^19–23^. While less is known about IMs, they are thought to provide a secondary response supporting both pathogen clearance and the resolution of inflammation^17, 24–28^. Depending on the nature of the challenge, lung macrophages must adapt their phenotype, accordingly, often referred to as ‘polarisation’ these functional states are broadly split into M1 pro-inflammatory and M2 anti-inflammatory macrophages although this nomenclature fails to capture the range of phenotypes that occur in macrophages^29, 30^. Metabolic alterations underpin macrophage function and, while these processes are well defined *in vitro* using model bone marrow derived macrophages (BMDMs), less is known about tissue macrophage polarisation. In response to LPS *in vivo*, AMs fail to elevate extracellular acidification and do not upregulate glycolytic pathways, instead relying on OXPHOS for cytokine production^31–33^. Despite this fundamental metabolic difference, in direct contrast to the glycolytic profile of BMDMs responding to LPS *in vitro*, metabolic characterisation of AMs following exposure to IL-4 has been overlooked *in vivo*. While it has been suggested that IMs may use mitochondrial metabolism to reduce LPS-induced lung injury, their metabolic requirements for polarisation remain poorly described. During murine *Mycobacterium tuberculosis* infection^25, 34^, AM and IM metabolism was directly compared, with the authors identifying that AMs become more M2 like (enhancing fatty acid oxidation (FAO)) while IMs were more glycolytic (supporting an M1 like metabolic profile)^17, 18^. However, it remains to be determined if a similar metabolic distinction is observed in the lung macrophage response to polarising cytokines, or in other inflammatory contexts.

One of the key challenges facing metabolic studies of tissue macrophages is removal from the complex tissue environment can disrupt expression of macrophage defining markers^15, 35–39^. In order to rapidly profile the metabolic state of lung macrophages with minimal loss of *in vivo* phenotype, we applied novel flow cytometry techniques SCENITH and Met-Flow, as well as RNA sequencing (RNAseq) of isolated macrophages following their polarisation *in vivo*. Our data has previously suggested that AMs may be reliant upon glucose metabolism in their response to IL-4 *in vitro*^15^. We now extend this work to reveal that naïve AMs have a high reliance upon glucose metabolism that was evident following *in vivo* polarisation with either IL-4 or LPS. RNA sequencing identified that AMs responding to IL-4 upregulated metabolic features of M2 polarisation (like OXPHOS, N-glycan biosynthesis and arginine metabolism). Flow cytometric analysis further supported these results, identifying metabolic alterations in AMs stimulated with IL-4 *in vitro* that were not observed in IMs. Additionally, both RNAseq and flow cytometric analysis showed that AMs were metabolically less responsive to LPS than BMDMs, failing to adopt a glycolytic phenotype or enrich any metabolic pathways at a transcriptional level in response to LPS *in vivo*. Contrastingly, although IMs failed to alter their metabolic profile in response to IL-4 *in vivo*, they acquired a glycolytic phenotype characteristic of M1 macrophages following *in vivo* LPS exposure. Finally, we show that glycolysis was enriched in AMs responding to IL-4 *in vivo* and reveal that their M2 polarisation was highly sensitive to glucose inhibition *ex vivo*. Together, we demonstrate that lung macrophage subsets display distinct metabolic profiles following classical and alternative activation *in vivo,* opening possibilities for selective metabolic inhibition of lung macrophage responses to modulate disease outcomes.

## Results

### Immune cells in the murine lung display distinct metabolic phenotypes

In recent years, novel flow cytometry techniques have been developed to enable the metabolic profiling of heterogenous and rare cell populations at a single cell level, with a resolution that would be challenging using Seahorse, long the gold standard for metabolic studies^40–42^. Single Cell ENergetIc metabolism by profiling Translation inHibition (SCENITH) allows for the quantification of global metabolic functions by measuring rates of translation^43^, while MET-flow directly measures the expression of a range of transporters and enzymes central to glycolysis and OXPHOS^44^. We first used both approaches to enable a direct metabolic comparison between immune cells in the naïve murine lung with a level of granularity previously not possible by Seahorse.

SCENITH relies on protein translation as a measure of global metabolic activity, with ATP levels correlating with protein synthesis rates^43^. The use of metabolic inhibitors targeting glucose and mitochondrial metabolism differentially impacts ATP production in different cell types, measured by SCENITH as a reduction in protein synthesis, with these changes in protein synthesis used to calculate cellular metabolic dependencies. Using this approach, we found that immune cells isolated from murine lung tissue with minimal exposure to culture medium underwent active protein translation under homeostasis that was sensitive to metabolic inhibition, enabling analysis by SCENITH (Fig. 1A). Despite the low glucose environment of the airways, AMs displayed a high glucose dependence, a phenotype shared by IMs and conventional dendritic cell (DC) subsets (cDC1s and cDC2s)^45^, suggesting a role for glucose metabolism in pulmonary macrophages and DCs (Fig. 1B). Both lung macrophage subsets and cDC1s displayed a high mitochondrial dependence, significantly greater than cDC2s, (Fig. 1C). Lymphocytes (B cells and CD4^+^, CD4^-^ and regulatory T cell subsets) showed high mitochondrial reliance, while neutrophils appeared to be completely reliant on glycolysis.

**Fig 1.**
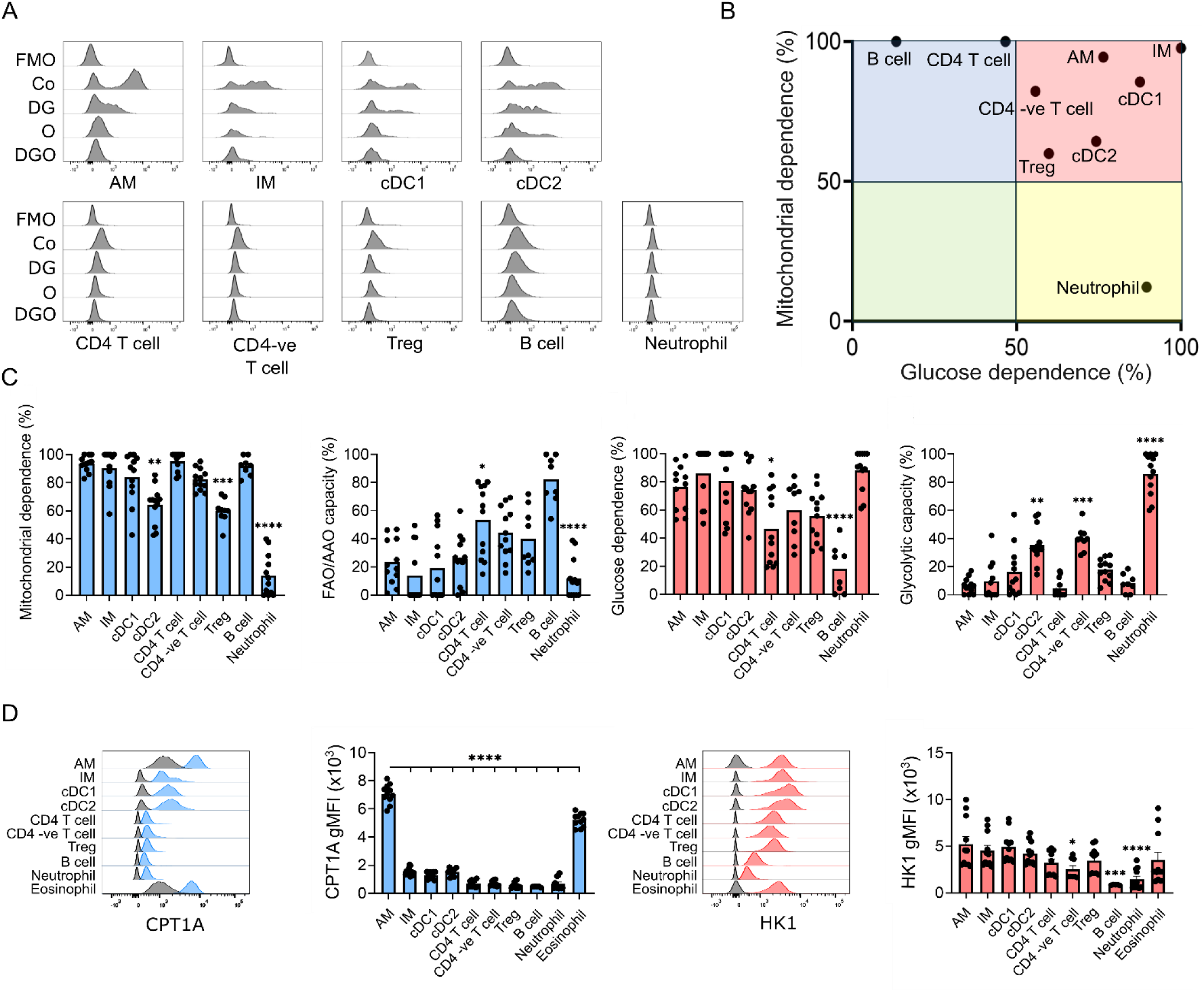
Metabolic analysis of macrophages and dendritic cells in the murine lung. Lungs from naïve C57BL/6 mice were processed to a single cell suspension prior to SCENITH analysis. **A** Representative histograms of puromycin levels in immune populations either untreated or treated with 2DG, O or 2DG and O. **B** Summary scatter plot showing mean mitochondrial and glucose dependencies. **C** Individual data points for glucose and mitochondrial dependencies as well as FAO/AAO and glycolytic capacities calculated from changes in puromycin levels upon addition of metabolic inhibitors. **D** Flow cytometric quantification of the expression levels of metabolic enzymes CPT1a and HK1 shown as bar charts and representative histograms. Data pooled from three independent experiments, each data point is an independent murine sample n= >10. Statistical analysis performed prior to data handling using a one-way ANOVA with *post hoc* Tukey tests compared to the AM group where **p <0.01, ***P <0.001 and ****p <0.0001.

As a complementary approach to SCENITH, to assess the role of lipid metabolism in AMs and glycolysis in IMs, rate limiting enzymes in FAO (CPT1A) and glycolysis (HK1) were quantified by MET-flow. The level of CPT1A was significantly higher in AMs than their IM counterparts, while HK1 expression was similar in both lung macrophage subsets, despite environmental differences in glucose availability in the airways versus tissues (Fig. 1D).

### Lung macrophage subsets display different metabolic responses to polarising stimuli *in vivo*

Following our interrogation of metabolic profiles of lung macrophage subsets in naïve mice, we next assessed if SCENITH could distinguish metabolic phenotypes of lung macrophages in different activation states. Contrasting the high metabolic dependencies we observed in *ex vivo* lung macrophages (Fig. 1), we found that naive (unstimulated) BMDMs were metabolically flexible (Fig. 2A). When we polarised BMDMs *in vitro* with IL-4 or LPS/IFN-γ, we found that IL-4 did not significantly alter the metabolic phenotype sustaining metabolic flexibility suggested by Seahorse analysis^46^, unlike LPS/IFN-γ which induced elevated glucose dependence with a concomitant loss of mitochondrial dependence supporting previous SCENITH analysis^47^ (Fig. 2A). Together, these data demonstrated that SCENITH was able to capture the highly glycolytic phenotype of M1 BMDMs established by Seahorse analysis *in vitro*, consistent with previous reports^46, 48^. After establishing SCENITH as a tool to characterise BMDM metabolic alterations driven by polarising stimuli *in vitro*, we wanted to determine if airway macrophages undergo similar metabolic alterations following polarisation *in vivo*. Intranasal delivery of PBS, LPS or IL-4c (a complex of recombinant IL-4 and an anti-IL-4 monoclonal antibody^49^ used to sustain the half life of IL-4 *in vivo*), did not significantly alter AM metabolic dependencies (Fig. 2B). However, expression of mitochondrial (CPT1A and ATP5a) and a glucose processing enzyme (HK1) was elevated in AMs following IL-4c delivery, suggesting increased fatty acid and glucose flux (Fig. 2C). Like AMs, IL-4c *in vivo* had no significant effect on IM metabolism, but LPS-driven polarisation induced a glycolytic phenotype characteristic of M1 macrophages we have defined *in vitro* (Fig. 2A). Despite this metabolic re-wiring, we did not observe significant changes in levels of glucose or mitochondrial enzymes in LPS stimulated IMs (Fig. 2C).

**Fig. 2.**
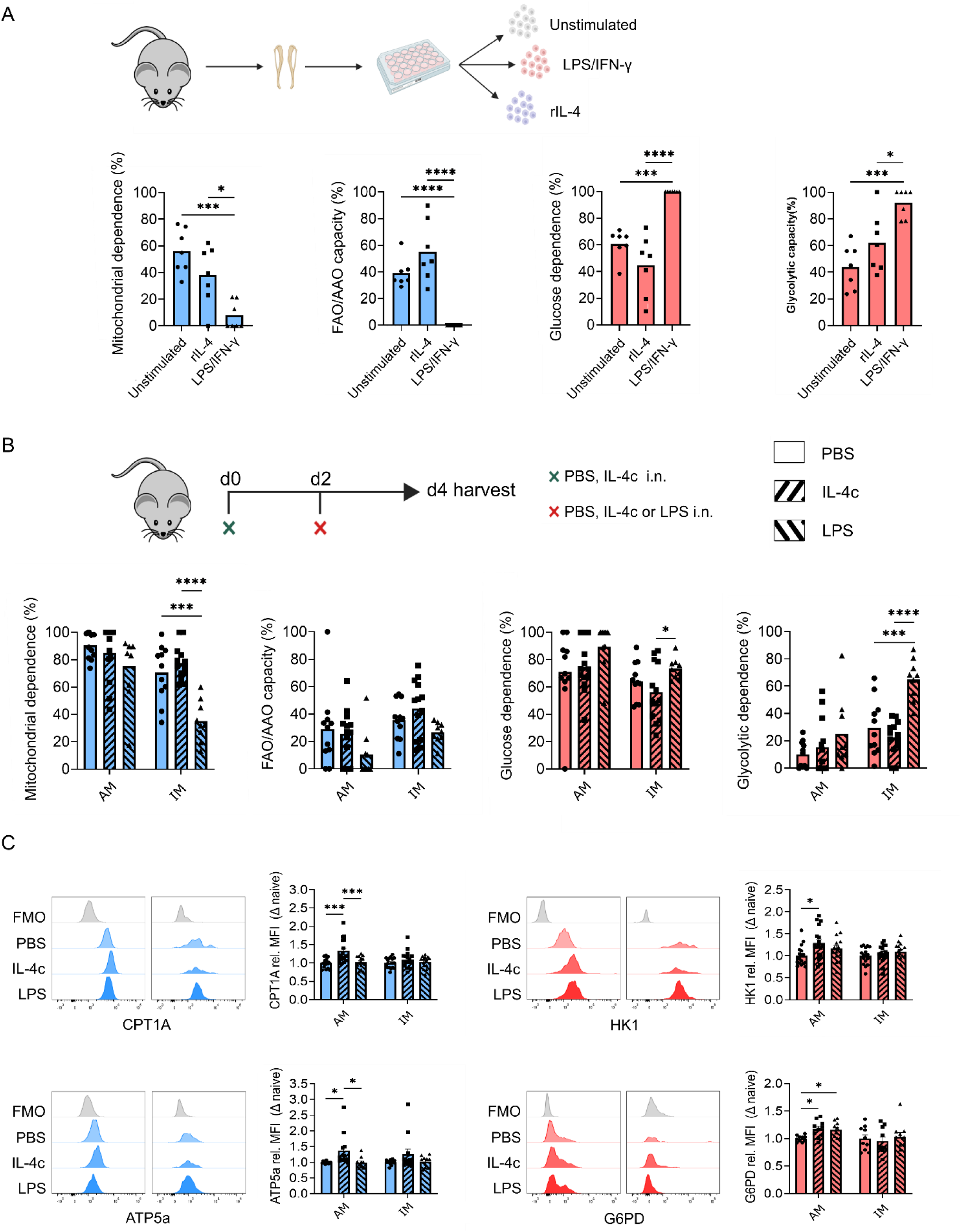
AMs and IMs are metabolically more responsive to IL-4 and LPS respectively. **A** BMDMs were left untreated or cultured with 20 ng/ml IL-4 or 20 ng/ml LPS and IFN-γ for 24 hours prior to SCENITH analysis. **B** Polarising cytokines were delivered intranasally, PBS or 0.05 µg IL-4c (days 0 and 2) or 0.15 µg LPS (day 2). Mice were harvested on day 4, lungs digested and SCENITH analysis performed on AMs. **C** Flow cytometric quantification of the expression levels of metabolic enzymes CPT1A, HK1 ATP5a and G6PD in polarised AMs shown as bar charts and representative histograms. SCENITH data pooled from two independent experiments, each data point is an independent murine sample n= 7. Each BMDM data point represents cells generated from a separate mouse. Statistical analysis performed prior to data handling using a one-way ANOVA with *post hoc* Tukey tests where *p <0.05, ***P <0.001 and ****p <0.0001.

### AMs are metabolically more responsive to IL-4c than LPS

To understand the metabolic response of tissue macrophages polarised *in vivo* in the lung environment, we next performed RNAseq on macrophages isolated *ex vivo* 24 hours after intranasal delivery of PBS, IL-4c or LPS. For this approach we focussed on AMs, as we could not undertake the same *ex vivo* RNAseq approach with IMs due to limiting cell numbers. Principal component analysis (PCA) revealed that *in vivo* exposure to IL-4c or LPS induced AMs to become distinct from PBS treated mice, significantly altering expression of 341 and 203 transcripts, respectively (Fig. 3A,B). Since 39 publicly available data sets were recently integrated to identify the most upregulated genes following polarisation of peritoneal macrophages, RAW264.7 and BMDMs with LPS and IL-4 (although ∼80% of data sets used BMDMs activated *in vitro*), we next determined if the same genes were upregulated by AMs after their polarisation *in vivo*^50^. IL-4c increased expression of M2 associated genes like *Chil3*, *Il31rα* and *Pdcd1lg2* (PD-L2) but, in keeping with our previous work^15^, failed to strongly induce *Retnlα* expression, a well-known marker of M2 polarised BMDMs (Fig. 3C)^15^. KEGG pathway analysis revealed that IL-4c upregulated genes associated with glycolysis and features of *in vitro* M2 activation, like OXPHOS, N-glycan biosynthesis and arginine metabolism (Fig. 3D)^51, 52^. AM expression of M1 associated genes like *Il6*, *Cxcl10* and *Ccl5* was increased by *in vivo* LPS exposure (Fig. 3C). However, KEGG analysis revealed that, differentially to IL-4c, *in vivo* LPS exposure failed to significantly upregulate metabolic pathways in AMs, while also identifying an increase in toll-like receptor (TLR), TNF and NF-κb signalling pathways which are known to mediate LPS driven M1 polarisation in BMDMs *in vitro*^51, 52^.

**Fig. 3.**
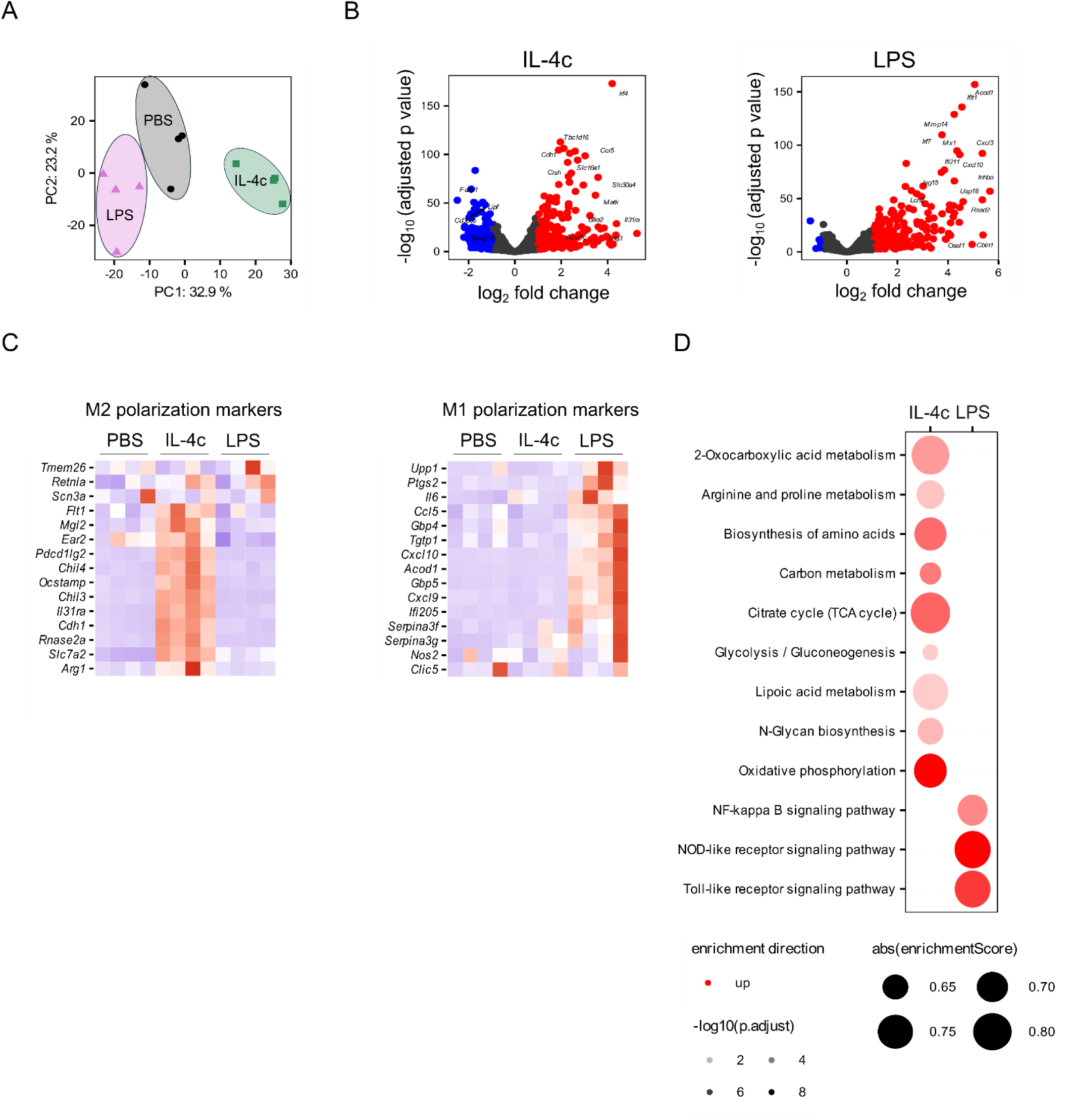
AMs are more metabolically responsive to IL-4c than LPS. C57BL/6 female mice received 40 µl of PBS, 0.05 µg IL-4c in 40 µl of PBS or 0.15 µg LPS in 40 µl PBS intranasally on d0. After 24 hours lungs were digested and sorted as described in (Supplementary Fig 2) before RNA extraction and bulk RNA sequencing. **A** PCA plot shows separate clustering of PBS, IL-4c and LPS treated macrophages. **B** mRNA expression profile represented as a volcano plot where significantly upregulated genes (red) and significantly downregulated genes (blue) are presented in IL-4c vs PBS and LPS Vs PBS. **C** Heat maps of differentially expressed genes between PBS, IL-4c and LPS of M2 and M1 associated genes. **D** KEGG pathway enrichment analysis selected for significantly enriched metabolic pathways and signalling pathways.

### Glucose metabolism is required for alternative activation *ex vivo*

Having identified that AMs upregulated both OXPHOS and glycolysis in response to IL-4 *in vivo* (Fig. 3D), we next used *ex vivo* inhibitor studies to dissect the metabolic requirements for AM production of type 2 mediators. BAL was harvested to isolate AMs with a ∼90% purity (supplementary figure 3) and to establish a functional link between AM metabolism and M2 activation, we examined the impact of the metabolic inhibitors 2DG (glucose metabolism) and etomoxir (FAO) on AM secretion of type 2 mediators (CCL17, CCL22 and YM1). Supporting the role for glucose metabolism in AMs polarised with IL-4 *in vivo* (Fig 2B), we found that AMs were dependent on glucose metabolism in response to IL-4, with inhibition significantly impairing production of these type 2 mediators following addition of 1 mM 2DG. Further, even a very low concentration of 2DG (0.2 mM) significantly inhibited AM secretion of CCL17 and CCL22, but not YM1 (Fig. 4A). Considering CCL17 and CCL22 are under control of phosphorylated Signal transducer and activator of transcriptions 6 (pSTAT6), but YM1 has both pSTAT6 dependent and independent mechanisms^53^, we next investigated whether the inhibitory effects of 2DG at low concentrations were pSTAT6 dependent. Glucose inhibition did not affect STAT6 phosphorylation, even at concentrations significantly higher than those needed to inhibit chemokine production (Fig. 4B). To further investigate metabolic requirements for AM polarisation, we assessed AM reliance on FAO and synthesis, previously linked to M2 activation and a known requirement for AM identity, for type 2 mediator secretion^54, 55^. Disruption of FAO and synthesis using the inhibitors etomoxir and C75 was found to impair M2 activation in a pSTAT6-dependent manner and inhibition of fatty acid synthesis but not oxidation impaired production of YM1 (Fig. 4C). Lastly, establishing the role of OXPHOS in M2 activation, the ATP synthase inhibitor oligomycin^56^ successfully prevented IL-4 induced upregulation of pSTAT6 in AMs (Fig. 4D).

**Fig 4.**
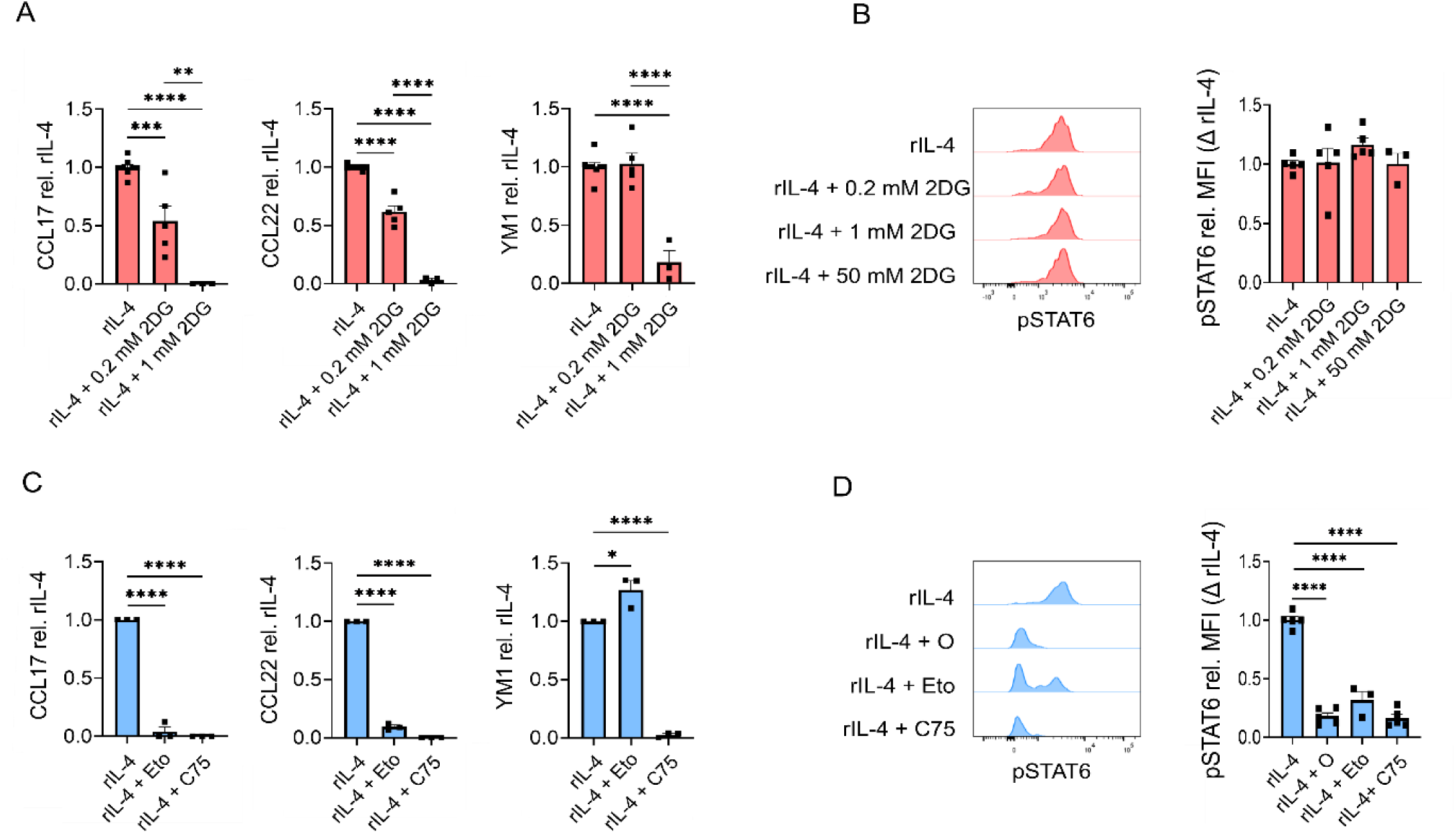
2DG impairs alternative activation independent of pSTAT6 signalling. BAL was harvested from naïve C57BL/6 female mice isolating AMs. AMs were allowed to adhere prior to addition of 20 ng/ml rIL-4 alone or rIL-4 + metabolic inhibitor. Type 2 molecules secreted by AMs was assessed by ELISA after addition of **A** 0.2 mM 2DG, 1mM 2DG **C** 200 µM etomoxir or 50 µM C75. Flow cytometric quantification of STAT6 phosphorylation following addition of **B** 0.2 mM 2DG, 1mM 2DG **D** 1 µM oligomycin, 200 µM etomoxir or 50 µM C75. Data pooled from at least two independent experiments, each data point is from 5-8 mice pooled to form a single biological replicate n= ≥3. Statistical analysis performed a one-way ANOVA with *post hoc* Tukey tests where *p <0.05, **p <0.01, ***P <0.001 and ****p <0.0001.

In summary, we identified that M2 polarisation of AMs is reliant upon both glycolytic and fatty acid metabolism, and that the inhibitory effect of 2DG on M2 activation is independent of pSTAT6 signalling.

## Materials and Methods

### Experimental animals

C57BL/6 mice (Envigo) were maintained under specific pathogen free (SPF) at the University of Manchester Biological Services Facility. All experiments were performed under the approval of project licenses granted by the Home Office U.K and conducted in accordance with local guidelines and with the United Kingdom Animals and Scientific Procedures Act of 1986. Age matched female mice aged 6 - 14 weeks were used in experiments and euthanised using a rising concentration of carbon dioxide (CO_2_). We acknowledge the study is limited to female mice and that sex differences can influence immunological and metabolic outcomes.

### Isolation of immune cells via lung digestion and BAL

Lungs were diced with a razor blade and incubated at 37°C for 40 minutes with 1.6U of TL liberase (Roche) and 160U DNase (Sigma) in H9269 (Sigma-Aldrich). Following digestion cold FACS buffer was used to stop digestion before lungs were crushed through a 70 μM filter to a single cell homogenate. RBC lysis buffer (Sigma) was used to lyse erythrocytes before cells were counted and used in downstream assays or processed for flow cytometry staining. BAL was obtained by washing the airway with 2% FCS and 2 mM EDTA (Sigma) in PBS.

### Intranasal delivery of IL-4c and LPS

Complexes of recombinant murine IL-4 (rIL-4) (PeproTech) and rat IgG1 anti-IL-4 monoclonal antibodies (IL-4 mAB) (BioXcell) were combined at a 1:5 ratio and referred to as IL-4 complex (IL-4c)^57^. IL-4c composed of 0.05 µg rIL-4 and 0.25 µg IL-4 mAB or 0.15 µg of lipopolysaccharide (LPS) (MERK) was delivered intranasally (i.n.) under anaesthetic (isoflurane) in 40 µl of sterile phosphate buffered saline (PBS) (Sigma-Aldrich). SCENITH and Met-Flow analysis were performed on day 4 following delivery of 40 µl PBS, 0.05 µg IL-4c on days 0 and 2 or 0.15 µg of LPS on day 2.

### Viability, surface and intracellular staining

Single cell suspension was washed with PBS and stained with Zombie ultraviolet (UV) dye (Biolegend) diluted 1:2000. Cells were washed with FACS buffer (PBS containing 2% FBS and 2 mM EDTA) to sequester unbound Zombie ultraviolet dye prior to blocking with 5 μg/ml FC block (αCD16/CD32; 2.4G2; Biolegend), surface staining was subsequently performed for 30 minutes at 4°C in FACS buffer. For intracellular staining, cells were fixed and permeabilised using ebioscience (ebio) fixation/permeabilization buffer for 45 minutes at 4°C. Cells were washed with 1X ebio permeabilization buffer (Thermofisher) before overnight staining. Stained cells were acquired using the 5 laser LSR fortessa using BD FACS Diva software and samples were analysed using FlowJo version 10.10 Software (Tree Star, Inc).

### SCENITH

Following BMDM harvest or lung tissue digestion cells were counted and 0.5X10^6^ cells seeded into each group: control (no inhibitor), 2-Deoxy-D-glucose (DG) (glucose inhibition), oligomycin (O) (mitochondrial inhibition) and DGO (both glucose and mitochondrial inhibition). Cells were incubated for 15 minutes at 37°C 5% CO_2_ before inhibitors were added for a further 15 minutes, the O in 2DGO which was added for the final 5 minutes of incubation. Puromycin was added for a further 30 minute incubation at 37°C 5% CO_2_ before ice cold PBS was immediately added to slow cellular metabolism. Cells were washed before proceeding to flow staining protocol previously described. Puromycin staining was performed using anti-puromycin AF647 clone R474 (now available at GammaOmics) or 2A4.

Geometric mean fluorescent intensity (gMFI) was used in dependency calculations listed below:

**Co = Control**

**DG = 2-Deoxy-D-glucose**

**O = Oligomycin**

**DGO = 2-Deoxy-D-glucose + Oligomycin**

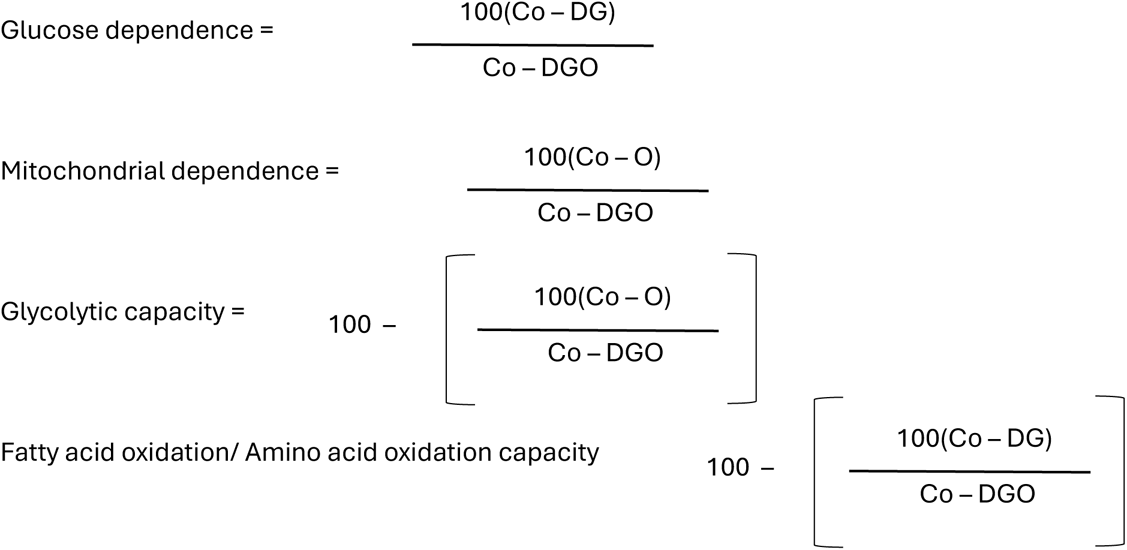

### Generation of BMDMs and *ex vivo* culture of AMs

Bone marrow cells were extracted from murine bones and cultured in complete RPMI 1640 (containing 10% FBS, 1% PenStrep, 1% l-glutamine, Sigma) in the presence of 20 ng/ml M-CSF (Peprotech) for 7 days. Culture medium was replaced ± IL-4 or LPS (both 20 ng/ml) for 24 hours before proceeding to SCENITH protocol. AMs were cultured in complete RPMI 1640 for 48 hours at 37°C 5% CO_2_ in the presence of IL-4 (20 ng/ml) ± 0.2 mM or 1 mM 2DG, 200 µM etomoxir or 50 µM C75 (all Sigma).

### pSTAT6 intracellular staining

Following BAL isolation AMs were incubated for 15 minutes with either 0.2 mM, 1 mM or 50 mM 2DG, 200 µM etomoxir, 1µM oligomycin (Sigma) or 50 µM C75 prior to a 30 minute stimulation with 20 ng/ml IL-4 (Peprotech) before proceeding to flow cytometry staining. Live/dead staining was performed as previously described. Cells were fixed with 1% PFA for 10 minutes at room temperature in the dark. For permeabilization Perm buffer III (BD) was added to cells for 30 minutes on ice, before blocking and surface staining as previously described. pSTAT6 staining was performed using anti-pSTAT6 PE (clone pY641) and the isotype control IgG1 R-PE Conjugate (Invitrogen). Cells were incubated with respective antibodies for 45 minutes at 4°C and washed prior to acquisition as previously described.

### Alveolar macrophage sorting

Lungs were processed as previously described and samples were both blocked and surface stained with Fc block (Biolegend), BV510 CD45, APC MerTK, PE CD64, BV711 CD11b and PE/CF594 Siglec-F in 500 μl of lung buffer for 30 minutes at 4°C. Following staining cells were washed prior to adding 0.25 µg/ml of the viability dye 4′,6-diamidino-2-phenylindole (DAPI) (SigmaAldrich) was added to enable the sorting of live cells. Cells were fluorescence activated cell sorting (FACS) on the BD Influx (BD Biosciences) at a pressure of 7.5 psi using a 140 µm nozzle to sustain the health of myeloid cells.

### RNA extraction, library construction and analysis

To generate samples for sequencing, mice were intranasally exposed to PBS, IL-4c (0.05 µg) or LPS (0.15 µg) 24 hours prior to harvest. Five mice were pooled to generate each biological replicate and four biological replicates from AM PBS, AM IL-4c and AM LPS groups were used in sequencing. After FACS sorting as described above, cells were lysed using RLT buffer (Qiagen) and RNA was extracted using RNeasy micro kits (Qiagen) according to manufacturers instructions. RNA quality and integrity was assessed using TapeStation (Agilent) before isolated RNA was shipped on dry ice to Novogene (Cambridge, UK) for library preparation and sequencing. mRNA was isolated using magnetic beads bound with poly-T oligos and then fragmented. First strand cDNA synthesis was performed using random hexamer primers, then second strand synthesis was performed with dTTPs. Strands were end-repaired and A-tailed, adapters were ligated, strands were then selected for size, amplified and purified. Libraries were pooled and sequenced using a NovaSeq X Plus PE150.

Raw reads were processed using in-house scripts at Novogene (Cambridge, UK). Hisat2 (v2.0.5) was used to align clean reads to the GRCm38.p6 Mus musculus reference genome. FeatureCounts (v1.5.0-p3) was used to generate read counts.

Differential expression analysis was performed using DESeq2 (v1.44.0) for R. Log fold change shrinkage was performed using the ashr package. Genes were considered significantly differentially expressed when the adjusted p value (padj) < 0.05 and the absolute log2(FoldChange) > 1. Dimensionality reduction was performed by Principal Components Analysis (PCA) on all genes using the ‘prcomp’ function. Read counts were transformed using the ‘varianceStabilizingTransformation’ function prior to PCA analysis. Gene-set enrichment analysis of Kyoto Encyclopedia of Genes and Genomes (KEGG) terms was performed using the clusterProfiler package. Nonsensical KEGG terms were manually filtered out prior to plotting. For generating heat maps normalized read counts were scaled across samples for each gene using the ‘scale’ function.

### ELISA

ELISAs to detect CCL17, CCL22 and YM1 (R&D SYSTEMS) were performed from AM culture supernatants according to manufacturers instructions.

### Statistical analysis

Statistical analyses were performed using GraphPad Prism 9 or JMP software. As described in figure legends, experimental groups were compared using a one-way analysis of variance (ANOVA) with Tukey test. Significance for all statistical tests was shown in figures as P < 0.05 (*), P < 0.01 (**), P < 0.001 (***), P < 0.0001 (****).

## Discussion

We have revealed that, despite residing in a low glucose environment, AMs invest in glucose processing pathways required to sustain protein translation (Fig. 1). We have demonstrated that IMs fail to alter their metabolic phenotype during ‘alternative’ activation in response to IL-4c *in vivo,* but undergo glycolytic re-wiring in response to intranasal LPS. Interestingly, AMs were metabolically more responsive to IL-4c than LPS *in vivo*, upregulating features of alternative activation at both transcriptional and protein levels (Figs. 2 and 3). Mechanistically, we have found that glucose metabolism is a feature of AMs following both alternative/M2 and classical/M1 activation *in vivo* and revealed that metabolic inhibition of glucose metabolism impairs *ex vivo* polarisation of AMs independent of pSTAT6, a master regulator of alternative activation (Fig. 4). Thus, lung macrophages rely upon glucose metabolism during their polarisation but display distinct metabolic responses to IL-4c vs LPS.

Our SCENITH analysis supports previous Seahorse studies that have proposed a role for mitochondrial metabolism in lymphocytes (B cells, CD4^+^ T cells and CD4^-^ T cells)^58–60^ and glycolysis in neutrophils^61^, as well as previously reported metabolic distinctions between DC subsets^45, 62^. However, current understanding of lung macrophage metabolism remains limited, particularly in IMs where studies rely on inflammation-induced expansion to enable metabolic analysis^17, 25^. Comparisons between AM and IM metabolism have therefore been restricted to diseased contexts, while baseline metabolic differences in these two distinct lung macrophage populations have not yet been clearly defined in naïve settings. Taking advantage of the ability of SCENITH to metabolically profile low cell numbers and rare immune populations^43^, we directly addressed this gap in the understanding. This revealed that AMs and IMs isolated from naive mice were simultaneously dependent upon both mitochondrial and glucose metabolism showing limited metabolic flexibility, a surprising finding considering BMDMs are typically associated with a high degree of metabolic plasticity *in vitro*^46^ and given the very low glucose availability in the lung airways^63^. Supportive of our results, naïve AMs have previously been shown to display a lipid metabolic programme, characterised by high levels of OXPHOS, as determined by bulk RNAseq and Seahorse analysis *ex vivo*^15, 64^. Consistent with a role for lipid metabolism in AMs, we found CPT1A expression was much higher in AMs than IMs, supporting the proposed metabolic adaptation of AMs to the lipid rich airways^65^. Apart from the indication folic acid might be important in IMs, we could not find reports detailing fundamental metabolic requirements of IMs in naive settings, with literature often relying on inflammation induced expansion of the IM pool prior to metabolic analysis. For example, it has been established that, in the context of *Mycobacterium tuberculosis* infection (MtB), IMs have some capacity for FAO^17^ and display higher rates of glycolysis than AMs^18^. Seahorse analysis revealed naïve AMs display a blunted rate of glycolysis^15^ and it has been suggested the acidification observed, used to calculate glycolytic rate is mitochondrially derived carbonic acid, rather than lactate generated via glycolysis^34^. We used SCENITH to identify a high glucose dependence in naïve AMs, supporting more recent work identifying a substantial glycolytic rate in AMs with functional importance in pulmonary homeostasis (Fig. 1)^66^. Furthermore, we revealed the level of HK1, a key enzyme that commits glucose for metabolic processing, was equally expressed in both lung macrophage subsets, even though AMs have access to dramatically less glucose in the airways than IMs in the tissues^67^. Therefore, we have revealed that naïve AMs invest in and rely on both glucose and mitochondrial metabolism despite AMs residing in a low glucose environment and requiring a lipid metabolic programme for ATP synthesis^68^, while naïve IMs show a requirement for glucose and mitochondrial metabolism despite high glucose availability.

Before assessing the metabolic response of lung macrophages to polarising stimuli, we first established whether SCENITH could capture BMDM metabolic reprogramming during polarisation, as has previously been suggested based on Seahorse analyses. In response to pro-inflammatory stimuli (LPS/IFN-γ)BMDMs exhibit Warburg like metabolism, impairing mitochondrial metabolism and re-wire toward glycolysis^69^. Metabolic profiling by SCENITH confirmed these previous results, identifying decreased mitochondrial dependence along with increased glucose dependence and glycolytic capacity following LPS and IFN-γ stimulation (Fig. 2). Conversely to M1 polarisation, M2 activation induces a well-established increase in OXPHOS, as determined by Seahorse extracellular flux analysis^70^. However, in agreement with previous reports ^47^, we did not see an increase in mitochondrial dependence in M2 BMDMs relative to unstimulated BMDM. An underappreciated feature of alternative activation in BMDMs is an enhanced glycolytic reserve, enabling greater rates of glycolysis in the absence of mitochondrial respiration^71^. Therefore, despite reports of increased OXPHOS in M2 BMDMs as determined by Seahorse analysis^52^, the improved ability to switch to glycolysis may ultimately cause M2 macrophages to have a lower mitochondrial dependence, as determined by SCENITH. M1 BMDMs display high rates of glycolysis and impaired mitochondrial metabolism determined by Seahorse analysis^46^, the inability to use OXPHOS when glycolysis is impaired enables SCENITH to capture the glycolytic phenotype of M1 macrophages validating the ability of SCENITH to determine metabolic profiles of polarized macrophages.

After establishing that SCENITH could capture BMDM metabolic phenotypes following their polarisation *in vitro,* we next applied this approach to determine the metabolic requirements for lung macrophage subsets responding to type 2 or type 1 polarising stimuli *in vivo*. Under homeostasis, AMs display some features of alternative activation, expressing high basal levels of some typical type 2 molecules (YM1 and CD206) and exhibiting an M2 like metabolic profile (characterised by high rates of lipid metabolism and OXPHOS)^15, 34, 42, 72, 73^. A limited number of studies have assessed lung macrophage metabolism following polarisation *in vivo*, these studies failed to interrogate IL-4c driven polarisation and have focused on transcription analysis in response to LPS^31–34^. We show the metabolic dependencies of lung macrophages were unaltered following IL-4c driven polarisation (Fig. 2), suggesting that the airway environment instructs AM metabolism toward an alternatively activated phenotype. Furthermore, our identification of increased levels of ATP5a and CPT1A in AMs, but not IMs, suggests AMs are more responsive to IL-4c and supports previous *in vitro* studies identifying OXPHOS and lipid oxidation as features of alternative activation^55, 74^. Our use of SCENITH demonstrated that IMs, but not AMs, acquired a glycolytic phenotype characteristic of previous reports of M1 macrophages *in vitro*, following LPS exposure *in vivo*^70^. These divergent metabolic responses are also observed following MtB infection where IMs appear to be more glycolytic than AMs, secreting higher quantities of lactate and upregulating HIF1α^18^, a master glycolytic regulator ^75^. The lack of glycolytic responsiveness in AMs is further supported by a model of innate training, where LPS fails to elevate ECAR 6 days following *in vivo* exposure^76^.

To investigate the acute metabolic response to AM polarisation *in vivo* we performed RNA sequencing analysis 24 hours after intranasal delivery of IL-4c and LPS. Although previous transcriptomic analysis has investigated AM metabolism following LPS polarisation *in vivo* ^32, 33^, the metabolic response of AM to IL-4c *in vivo* remains unclear. We previously addressed this issue by delivering two intraperitoneal doses of IL-4c before undertaking RNAseq on isolated AMs^15^. Subsequent analyses revealed that M2 markers were not significantly upregulated in AMs exposed twice to IL-4c *in vivo*, with very few differentially expressed genes (DEGs) identified^15^. Building upon and extending this previous work, we analysed AM phenotype and metabolic profile after intranasal delivery of a single dose of IL-4c. This approach identified over 300 DEGs in AMs exposed *in vivo* to IL-4c, including some typical M2 markers (Fig. 3). KEGG pathway enrichment analysis revealed OXPHOS, N-glycan biosynthesis and arginine metabolism were enriched following IL-4c delivery *in vivo*. Together, our protein and transcriptional data demonstrate that *in vivo* administration of IL-4c induces metabolic alterations in AMs but not IMs previously described in M2 BMDMs *in vitro*. Following LPS delivery we show glycolysis was not upregulated in AMs (Fig. 3), likely a reflection of the limited environmental glucose availability^64, 77, 78^. This finding supports previous transcriptional analysis suggesting glycolysis was not elevated 24 hours after LPS delivery in mouse^33^ and was actively downregulated in human following bronchial delivery^32^. Thus, we have confirmed that metabolic re-wiring towards glycolysis observed following LPS stimulation of BMDMs *in vitro* is not a feature of AMs responding to LPS *in vivo.* We also reveal that AM stimulation with LPS did not alter expression of key enzymes in mitochondrial and glucose metabolism at the protein level (Fig. 2C), or drive enrichment of any metabolic pathways at the transcriptional level (Fig. 3D). Together, these data suggest that airway residing AMs have impaired metabolic responsiveness to LPS *in vivo* that is not shared by IMs found in the bronchial interstitium. Considering that surfactant clearance is inherently linked to mitochondrial metabolism^64^, we speculate that conditioning by the airway environment prevents AMs from adopting an M1 like metabolic profile.

Given that AMs were metabolically more responsive to IL-4c than LPS *in vivo*, we isolated AMs from BAL to assess their metabolic requirements for alternative activation *ex vivo*. Despite the low rate of glycolysis evident in naïve AMs^15^ we found that a low concentration (0.2 mM) of 2DG effectively attenuated secretion of the type 2 molecules CCL17 and CCL22, but not YM1, by AMs cultured with IL-4 *ex vivo* (Fig. 4). It has previously been established that YM1 is expressed in naïve AMs in an IL-4Rα independent manner ^53^. To determine if IL-4Rα signalling was responsible for the differential impact of 2DG on these M2 products, we quantified levels of pSTAT6, a key transcription factor induced by IL-4rα engagement^79^. We found 2DG failed to impair IL-4 induced phosphorylation of STAT6 suggesting a different mechanism may be responsible for 2DG inhibition of CCL17 and CCL22. Unlike glucose metabolism, inhibition of lipid oxidation and synthesis impaired pSTAT6 formation and subsequent production of type 2 molecules CCL17 and CCL22 (Fig. 4), supporting suggested requirements for lipid metabolism in BMDMs responding to IL-4 *in vitro*^80, 81^. We also observed inhibition of fatty acid synthesis but not oxidation impaired the production of YM1 a surprising finding considering synthesis is not directly linked to ATP production.

However, the role of FAO in M2 activation is disputed, with recent studies suggesting that BMDM M2 polarisation is not influenced by CPT1A deficiency, and that reported pharmacological inhibition of the M2 genes *Retnla*, *Mgl2* and *Fabp4* by etomoxir may instead be due to off target depletion of the intracellular Acetyl-CoA pool^82, 83^. Thus, some caution should be taken when interpreting results generated using etomoxir to block metabolic processes.

In summary, we show AMs are metabolically more responsive to alternative activation and IMs are more responsive to classical activation *in vivo* with implications in the ability for each subset to enable effective clearance of respiratory pathogens or sustain wound healing responses. We suggest that this division of labour may be a general feature of macrophages in other organs, where the location of macrophage subsets may dictate the metabolic dependencies of their activation and function.

## Author contributions

A.S.M. and S.C. were responsible for conceptualisation and supervision, L.B., B.H., A.J.L.R., G.M. and A.P.A. conducted or enabled investigations, J.J.H. performed formal analysis, curation and writing - original draft. R.J.A. provided key assistance with the SCENITHTM methodology, data interpretation and discussion on SCENITH and data. A.S.M. was involved in funding acquisition. A.S.M. and S.C. were involved in writing (review and editing).

## Acknowledgements

The authors acknowledge the Lydia Becker Institute and MacDonald laboratory (University of Manchester) for scientific discussions and experimental help. We also acknowledge the University of Manchester flow cytometry facilities. J.J.H. was supported by a MRC iCASE studentship (with GSK) (MR/R015767/1), B.H. by a GSK studentship, MRC iCASE (MR/N013751/1) A.L.R G.M. by an EPSRC Programme Grant (EP/P00119X/1), R.J.A by ANR JCJC-Epic ZENITH N° ANR-20-CE14-0028 and A.S.M. by funding from the Lydia Becker Institute and the MRC (MR/W018748/1).

## Data and materials availability

Any data not explicitly evident in the paper will be available on reasonable request.

## References

1. Huang, X., Cao, M. & Xiao, Y. Alveolar macrophages in pulmonary alveolar proteinosis: origin, function, and therapeutic strategies. Front Immunol 14, (2023).

2. Wardle, E. N. Kupffer cells and their function. Liver 7, 63–75 (1987).

3. Li, Q. et al. Developmental Heterogeneity of Microglia and Brain Myeloid Cells Revealed by Deep Single-Cell RNA Sequencing. Neuron 101, 207–223.e10 (2019).

4. Zaman, R. & Epelman, S. Resident cardiac macrophages: Heterogeneity and function in health and disease. Immunity 55, 1549–1563 (2022).

5. Guilliams, M. & Svedberg, F. R. Does tissue imprinting restrict macrophage plasticity? Nat Immunol 22, 118–127 (2021).

6. Gibbings, S. L. et al. Transcriptome analysis highlights the conserved difference between embryonic and postnatal-derived alveolar macrophages. Blood 126, 1357–1366 (2015).

7. Whitsett, J. A., Wert, S. E. & Weaver, T. E. Alveolar surfactant homeostasis and the pathogenesis of pulmonary disease. Annu Rev Med 61, 105–119 (2010).

8. Sakai, M. et al. Liver-Derived Signals Sequentially Reprogram Myeloid Enhancers to Initiate and Maintain Kupffer Cell Identity. Immunity 51, 655–670.e8 (2019).

9. Laar, L. van de et al. Yolk Sac Macrophages, Fetal Liver, and Adult Monocytes Can Colonize an Empty Niche and Develop into Functional Tissue-Resident Macrophages. Immunity 44, 755–768 (2016).

10. Brennan, A. L. et al. Airway glucose concentrations and effect on growth of respiratory pathogens in cystic fibrosis. Journal of Cystic Fibrosis 6, 101–109 (2007).

11. Gill, S. K. et al. Increased airway glucose increases airway bacterial load in hyperglycaemia. Sci Rep 6, (2016).

12. Mallia, P. et al. Role of airway glucose in bacterial infections in patients with chronic obstructive pulmonary disease. Journal of Allergy and Clinical Immunology 142, 815–823.e6 (2018).

13. Olmeda, B., Martínez-Calle, M. & Pérez-Gil, J. Pulmonary surfactant metabolism in the alveolar airspace: Biogenesis, extracellular conversions, recycling. Annals of Anatomy 209, 78–92 (2017).

14. Batenburg, J. J., Longmore, W. J., Golde, L. M. G. Van & Daisy, E. A. THE SYNTHESIS OF PHOSPHAT∼YLCHOL∼E BY ADULT RAT LUNG ALVEOLAR TYPE II EPITHELIAL CELLS IN PRIMARY CULTURR. Biochimicu et Biophysics Ado 529, (1978).

15. Svedberg, F. R. et al. The lung environment controls alveolar macrophage metabolism and responsiveness in type 2 inflammation. Nat Immunol 20, 571–580 (2019).

16. Gibbings, S. L. et al. Three unique interstitial macrophages in the murine lung at steady state. Am J Respir Cell Mol Biol 57, 66–76 (2017).

17. Huang, L., Nazarova, E. V., Tan, S., Liu, Y. & Russell, D. G. Growth of Mycobacterium tuberculosis in vivo segregates with host macrophage metabolism and ontogeny. Journal of Experimental Medicine 215, 1135–1152 (2018).

18. Pisu, D., Huang, L., Grenier, J. K. & Russell, D. G. Dual RNA-Seq of Mtb-Infected Macrophages In Vivo Reveals Ontologically Distinct Host-Pathogen Interactions. Cell Rep 30, 335–350.e4 (2020).

19. Allard, B., Panariti, A. & Martin, J. G. Alveolar Macrophages in the Resolution of Inflammation, Tissue Repair, and Tolerance to Infection. Front Immunol 9, (2018).

20. Schneider, C. et al. Alveolar Macrophages Are Essential for Protection from Respiratory Failure and Associated Morbidity following Influenza Virus Infection. PLoS Pathog 10, (2014).

21. Xu, J. et al. Scavenger Receptor MARCO Orchestrates Early Defenses and Contributes to Fungal Containment during Cryptococcal Infection. The Journal of Immunology 198, 3548–3557 (2017).

22. Preston, J. A. et al. Alveolar macrophage apoptosis-associated bacterial killing helps prevent murine pneumonia. Am J Respir Crit Care Med 200, 84–97 (2019).

23. Tate, M. D., Pickett, D. L., Rooijen, N. van, Brooks, A. G. & Reading, P. C. Critical Role of Airway Macrophages in Modulating Disease Severity during Influenza Virus Infection of Mice. J Virol 84, 7569–7580 (2010).

24. Kawano, H. et al. IL-10-producing lung interstitial macrophages prevent neutrophilic asthma. Int Immunol 28, 489–501 (2016).

25. Zhou, B. et al. The angiocrine Rspondin3 instructs interstitial macrophage transition via metabolic–epigenetic reprogramming and resolves inflammatory injury. Nat Immunol 21, 1430–1443 (2020).

26. Bedoret, D. et al. Lung interstitial macrophages alter dendritic cell functions to prevent airway allergy in mice. Journal of Clinical Investigation 119, 3723–3738 (2009).

27. Guilliams, M. & Svedberg, F. R. Does tissue imprinting restrict macrophage plasticity? Nat Immunol 22, 118–127 (2021).

28. Ren, J. et al. Intranasal delivery of MSC-derived exosomes attenuates allergic asthma via expanding IL-10 producing lung interstitial macrophages in mice. Int Immunopharmacol 91, (2021).

29. Yunna, C., Mengru, H., Lei, W. & Weidong, C. Macrophage M1/M2 polarization. Eur J Pharmacol 877, (2020).

30. Murray, P. J. Macrophage Polarization. Annu Rev Physiol 79, 541–566 (2017).

31. Moore, P. K. et al. Single-cell RNA sequencing reveals unique monocyte-derived interstitial macrophage subsets during lipopolysaccharide-induced acute lung inflammation. Am J Physiol Lung Cell Mol Physiol 324, L536–L549 (2023).

32. Otto, N. A. et al. Metabolic adaptations of human alveolar macrophages upon activation by lipopolysaccharide in vivo. Scand J Immunol 93, (2021).

33. Xiao, Y. et al. GCH1 reduces LPS-induced alveolar macrophage polarization and inflammation by inhibition of ferroptosis. Inflammation Research 72, 1941–1955 (2023).

34. Woods, P. S. et al. Tissue-resident alveolar macrophages do not rely on glycolysis for LPS-induced inflammation. Am J Respir Cell Mol Biol 62, 243–255 (2020).

35. Gosselin, D. et al. Environment drives selection and function of enhancers controlling tissue-specific macrophage identities. Cell 159, 1327–1340 (2014).

36. Gosselin, D. et al. An environment-dependent transcriptional network specifies human microglia identity. Science (1979) 356, 1248–1259 (2017).

37. Mass, E. Delineating the origins, developmental programs and homeostatic functions of tissue-resident macrophages. Int Immunol 30, 493–501 (2018).

38. Gautiar, E. L. et al. Gene-expression profiles and transcriptional regulatory pathways that underlie the identity and diversity of mouse tissue macrophages. Nat Immunol 13, 1118– 1128 (2012).

39. Lavin, Y. et al. Tissue-resident macrophage enhancer landscapes are shaped by the local microenvironment. Cell 159, 1312–1326 (2014).

40. Bossche, J. Van den, Baardman, J. & Winther, M. P. J. de Metabolic characterization of polarized M1 and M2 bone marrow-derived macrophages using real-time extracellular flux analysis. Journal of Visualized Experiments 2015, (2015).

41. Grey, J. F. E., Townley, A. R., Everitt, N. M., Campbell-Ritchie, A. & Wheatley, S. P. A cost-effective, analytical method for measuring metabolic load of mitochondria. Metabol Open 4, 100020 (2019).

42. Wang, S., Liu, G., Li, Y. & Pan, Y. Metabolic Reprogramming Induces Macrophage Polarization in the Tumor Microenvironment. Front Immunol 13, (2022).

43. Argüello, R. J. et al. SCENITH: A Flow Cytometry-Based Method to Functionally Profile Energy Metabolism with Single-Cell Resolution. Cell Metab 32, 1063–1075.e7 (2020).

44. Ahl, P. J., et al. Met-Flow, a strategy for single-cell metabolic analysis highlights dynamic changes in immune subpopulations. Commun Biol 3, (2020).

45. Kratchmarov, R. et al. Metabolic control of cell fate bifurcations in a hematopoietic progenitor population. Immunol Cell Biol 96, 863–871 (2018).

46. Viola, A., Munari, F., Sánchez-Rodríguez, R., Scolaro, T. & Castegna, A. The metabolic signature of macrophage responses. Front Immunol 10, (2019).

47. Verberk, S. G. S., et al. An integrated toolbox to profile macrophage immunometabolism. Cell Reports Methods 2, (2022).

48. Kelly, B. & O’Neill, L. A. J. Metabolic reprogramming in macrophages and dendritic cells in innate immunity. Cell Res 25, 771–784 (2015).

49. Jenkins, S. J. et al. Local macrophage proliferation, rather than recruitment from the blood, is a signature of T H2 inflammation. Science (1979) 332, 1284–1288 (2011).

50. Colombo, G., Pessolano, E., Talmon, M., Genazzani, A. A. & Kunderfranco, P. Getting everyone to agree on gene signatures for murine macrophage polarization in vitro. PLoS One 19, (2024).

51. Rath, M., Müller, I., Kropf, P., Closs, E. I. & Munder, M. Metabolism via arginase or nitric oxide synthase: Two competing arginine pathways in macrophages. Front Immunol 5, (2014).

52. Jha, A. K. et al. Network integration of parallel metabolic and transcriptional data reveals metabolic modules that regulate macrophage polarization. Immunity 42, 419–430 (2015).

53. Sutherland, T. E. et al. Ym1 induces RELMα and rescues IL-4Rα deficiency in lung repair during nematode infection. PLoS Pathog 14, (2018).

54. Kieler, M., Hofmann, M. & Schabbauer, G. More than just protein building blocks: how amino acids and related metabolic pathways fuel macrophage polarization. FEBS Journal 288, 3694–3714 (2021).

55. Huang, S. C. C. et al. Cell-intrinsic lysosomal lipolysis is essential for alternative activation of macrophages. Nat Immunol 15, 846–855 (2014).

56. Shchepina, L. A. et al. Oligomycin, inhibitor of the F 0 part of H +-ATP-synthase, suppresses the TNF-induced apoptosis. doi:10.1038/sj.onc

57. Jenkins, S. J. et al. Local macrophage proliferation, rather than recruitment from the blood, is a signature of T H2 inflammation. Science (1979) 332, 1284–1288 (2011).

58. Weisel, F. J. et al. Germinal center B cells selectively oxidize fatty acids for energy while conducting minimal glycolysis. Nat Immunol 21, 331–342 (2020).

59. Zhang, L. & Romero, P. Metabolic Control of CD8+ T Cell Fate Decisions and Antitumor Immunity. Trends Mol Med 24, 30–48 (2018).

60. Yang, W., Yu, T. & Cong, Y. CD4+T cell metabolism, gut microbiota, and autoimmune diseases: Implication in precision medicine of autoimmune diseases. Precis Clin Med 5, (2022).

61 Jeon, J. H., Hong, C. W., Kim, E. Y. & Lee, J. M. Current understanding on the metabolism of neutrophils. Immune Netw 20, 1–13 (2020).

61. Du, X. et al. Hippo/Mst signalling couples metabolic state and immune function of CD8α+ dendritic cells. Nature 558, 141–145 (2018).

62. Garnett, J. P., Baker, E. H. & Baines, D. L. Sweet talk: Insights into the nature and importance of glucose transport in lung epithelium. European Respiratory Journal 40, 1269–1276 (2012).

63. Schneider, C. et al. Induction of the nuclear receptor PPAR-γ 3 by the cytokine GM-CSF is critical for the differentiation of fetal monocytes into alveolar macrophages. Nat Immunol 15, 1026–1037 (2014).

64. Marelli, G. et al. Lipid-loaded macrophages as new therapeutic target in cancer. J Immunother Cancer 10, (2022).

65. Albers, G. J. et al. Airway macrophage glycolysis controls lung homeostasis and responses to aeroallergen. Mucosal Immunol (2024).doi:10.1016/j.mucimm.2024.10.002

66. Baker, E. H. et al. Hyperglycemia and cystic fibrosis alter respiratory fluid glucose concentrations estimated by breath condensate analysis. J Appl Physiol 102, (1969).

67. Cloonan, S. M. “KEAP”ing Alveolar Macrophage Mitochondria Content in Chronic Obstructive Pulmonary Disease. Am J Respir Crit Care Med 207, 962–964 (2023).

68. Hard, G. C. SOME BIOCHEMICAL ASPECTS OF THE IMMUNE MACROPHAGE. Br. J. exp. Path (1970).

69. Runtsch, M. C. et al. Itaconate and itaconate derivatives target JAK1 to suppress alternative activation of macrophages. Cell Metab 34, 487–501.e8 (2022).

70. Mookerjee, S. A., Nicholls, D. G. & Brand, M. D. Determining maximum glycolytic capacity using extracellular flux measurements. PLoS One 11, (2016).

71. Remmerie, A. & Scott, C. L. Macrophages and lipid metabolism. Cell Immunol 330, 27– 42 (2018).

72. Wculek, S. K. et al. Oxidative phosphorylation selectively orchestrates tissue macrophage homeostasis. Immunity 56, 516–530.e9 (2023).

73. Vats, D. et al. Oxidative metabolism and PGC-1β attenuate macrophage-mediated inflammation.

74. Obach, M. et al. 6-Phosphofructo-2-kinase (pfkfb3) gene promoter contains hypoxia-inducible factor-1 binding sites necessary for transactivation in response to hypoxia. Journal of Biological Chemistry 279, 53562–53570 (2004).

75. Zahalka, S. et al. Trained immunity of alveolar macrophages requires metabolic rewiring and type 1 interferon signaling. Mucosal Immunol 15, 896–907 (2022).

76. Baker, A. D. et al. Targeted PPARγ deficiency in alveolar macrophages disrupts surfactant catabolism. J Lipid Res 51, 1325–1331 (2010).

77. Koo, S. jie & Garg, N. J. Metabolic programming of macrophage functions and pathogens control. Redox Biol 24, (2019).

78. Yu, T. et al. Modulation of M2 macrophage polarization by the crosstalk between Stat6 and Trim24. Nat Commun 10, (2019).

79. Covarrubias, A. J. et al. Akt-mTORC1 signaling regulates Acly to integrate metabolic input to control of macrophage activation. (2016).doi:10.7554/eLife.11612.001

80. Bidault, G. et al. SREBP1-induced fatty acid synthesis depletes macrophages antioxidant defences to promote their alternative activation. Nat Metab 3, 1150–1162 (2021).

81. Covarrubias, A. J. et al. Akt-mTORC1 signaling regulates Acly to integrate metabolic input to control of macrophage activation. (2016).doi:10.7554/eLife.11612.001

82. Divakaruni, A. S. et al. Etomoxir Inhibits Macrophage Polarization by Disrupting CoA Homeostasis. Cell Metab 28, 490–503.e7 (2018).

