## Supplemental Figure 1 for "Pulmonary macrophage subsets display distinct metabolic responses to polarising stimuli *in vivo*"

**Supplementary Figure 1 flow cytometry gating strategy for identification of immune populations in the naïve lung for SCENITH and Met-Flow analysis**

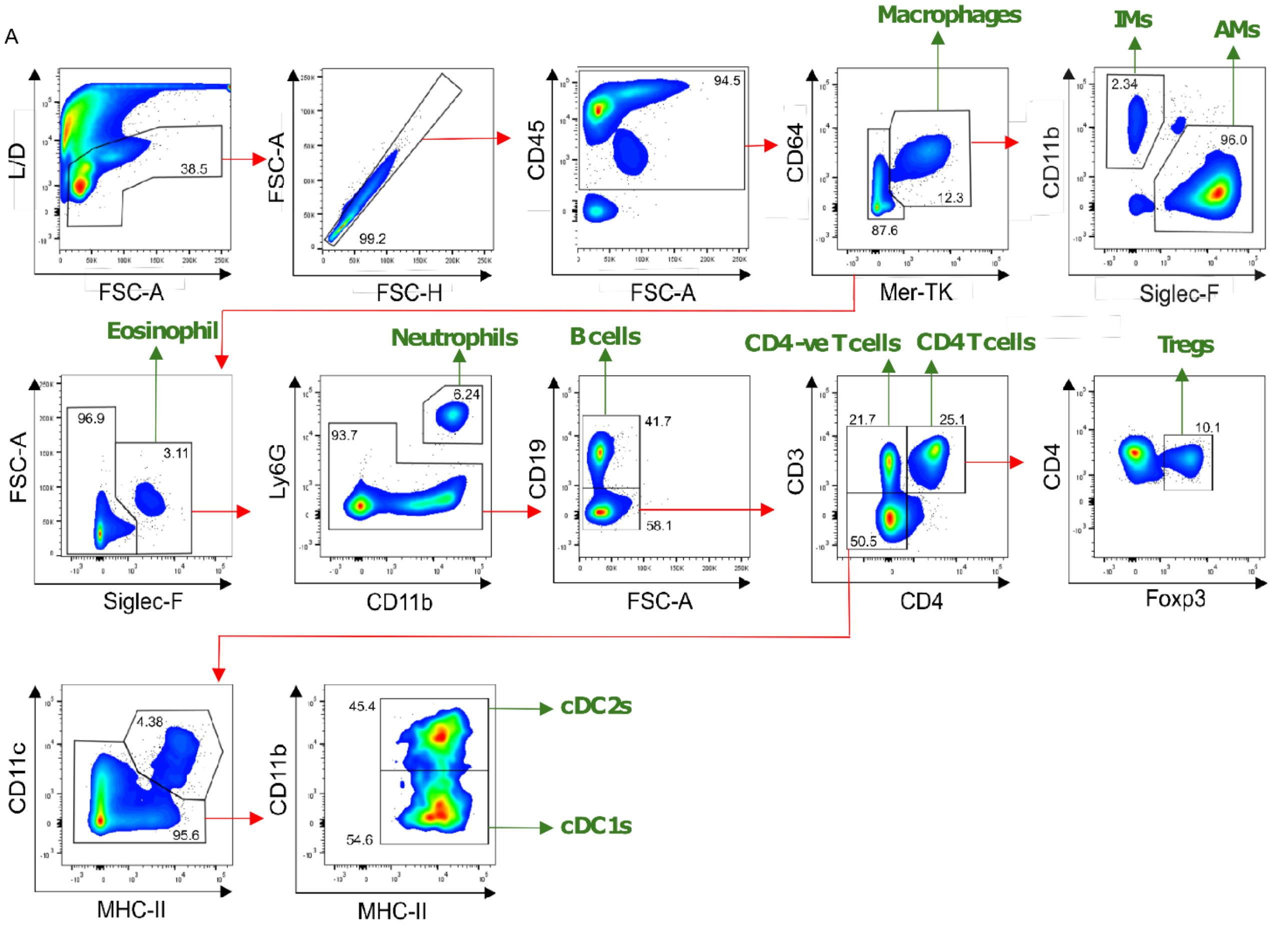


(**A**) Representative flow cytometry plots showing gating for the identification of cells isolated from naïve lung tissue, all populations identified were CD45+ identifying IM(MerTK+CD64+CD11b+Siglec-F-), AM (MerTK+CD64+CD11b-Siglec-F+), eosinophil (MerTK-CD64-Siglec-F+), neutrophils (MerTK-CD64- Siglec-F-Ly6G+CD11b+), B cells (MerTK-CD64-Siglec-F-Ly6G-CD11b-CD19+), CD4 -ve T cells (MerTK-CD64- Siglec-F-Ly6G-CD11b-CD19-CD3+CD4-), CD4 T cells (MerTK-CD64- Siglec-F-Ly6G-CD11b-CD19-CD3+CD4+), Tregs (MerTK-CD64- Siglec-F-Ly6G-CD11b-CD19-CD3+CD4+FOXP3+), cDC1s (MerTK-CD64- Siglec-F-Ly6G-CD11b-CD19-CD3-CD4-CD11b-MHC-II+), cDC2s (MerTK-CD64- Siglec-F-Ly6G-CD11b-CD19-CD3-CD4-CD11b+MHC-II+).
