## Supplemental Figure 2 for "Pulmonary macrophage subsets display distinct metabolic responses to polarising stimuli *in vivo*"

**Supplementary Figure 2 FACS sorting of alveolar macrophages from the murine lung following PBS, IL-4c or LPS treatment**

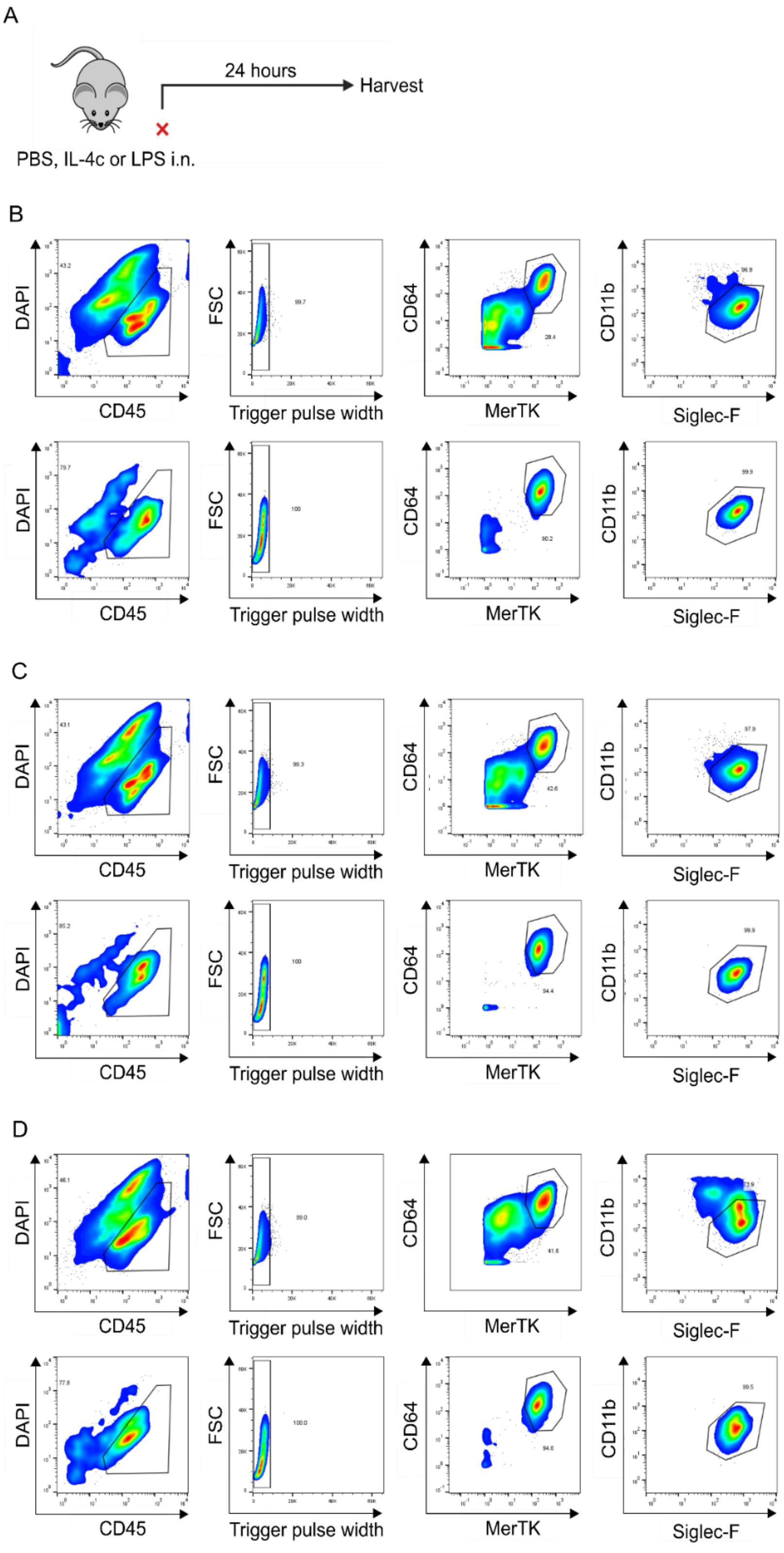

C57BL/6 female mice received 40 µl of PBS, (**A**) 0.05 µg IL-4c in 40 µl of PBS or 0.15 µg LPS in 40 µl intranasally on d0. Lungs were harvested after 24 hours, digested and stained with DAPI, CD45, CD64, MerTK, CD11b and Siglec-F. Gating strategy for alveolar macrophage sorting and sorted cells were run back through the sorter to determine purity following (**B**) PBS (**C**) IL-4c and (**D**) LPS. Gate frequencies show % of parent.
