## Supplemental Figure 3 for "Pulmonary macrophage subsets display distinct metabolic responses to polarising stimuli *in vivo*"

**Supplementary Figure 3 flow cytometry gating strategy for identification alveolar macrophages isolated by BAL from naïve lungs**

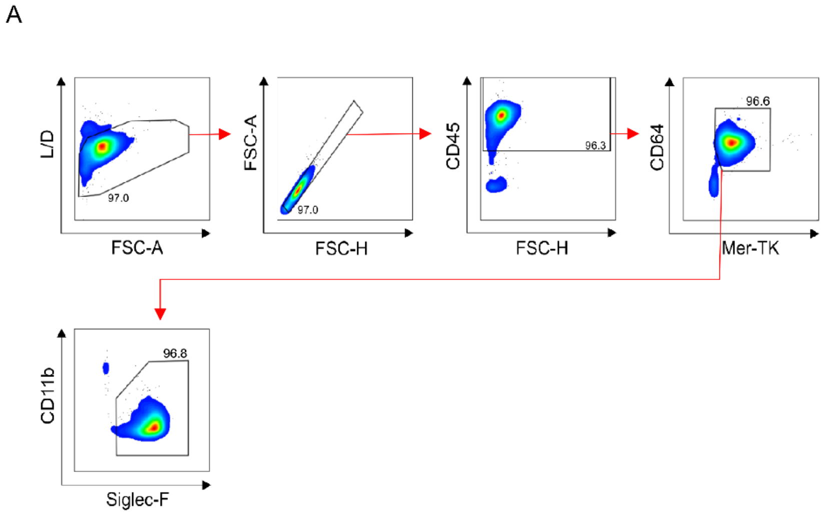


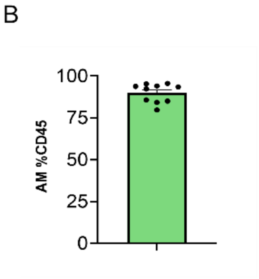


(**A**) Representative flow cytometry plots showing gating for the identification of AMs (CD45+MerTK+CD64+CD11b-Siglec-F+) isolated from BAL for ex vivo culture studies. (**B**) Purity of isolated AMs as %CD45+ cells isolated mean ± SEM, , each data point is an independent murine sample n= 10
